# Implementation of Bayesian Multiple Comparison Correction in the Second-Level Analysis of fMRI Data: With Pilot Analyses of Simulation and Real fMRI Datasets Based on Voxelwise Inference

**DOI:** 10.1101/855460

**Authors:** Hyemin Han

**Affiliations:** University of Alabama

**Author notes:** Author Note: The author thank Joonsuk Park and Ian McDonough for their comments on an earlier version of the manuscript.

## Abstract

We developed and tested Bayesian multiple comparison correction method for Bayesian voxelwise second-level fMRI analysis with R. The performance of the developed method was tested with simulation and real image datasets. First, we compared false alarm and hit rates, which were used as proxies for selectivity and sensitivity, respectively, between Bayesian and classical inference were conducted. For the comparison, we created simulated images, added noise to the created images, and analyzed the noise-added images while applying Bayesian and classical multiple comparison correction methods. Second, we analyzed five real image datasets to examine how our Bayesian method worked in realistic settings. When the performance assessment was conducted, Bayesian correction method demonstrated good sensitivity (hit rate ≥ 75%) and acceptable selectivity (false alarm rate < 10%) when N ≤ 8. Furthermore, Bayesian correction method showed better sensitivity compared with classical correction method while maintaining the aforementioned acceptable selectivity.

## 1 Introduction

We aimed at developing and testing a novel method for multiple comparison correction in second-level (group) fMRI analysis based on Bayesian statistics. In the field of neuroimaging, how to deal with the inflation of false positives due to multiple testing has been a fundamental methodological issue (Bennett, Miller, & Wolford, 2009). Given that usual fMRI analysis involves testing of more than ten to hundred thousand voxels, the possibility to encounter Type I error is likely to be significantly inflated once we adopt a widely used *p*-value threshold, *p* < .05, without any further treatment. To address this issue, researchers have developed various statistical methods (e.g., voxelwise and clusterwise familywise error rate correction and false-discovery rate correction) to adjust the aforementioned *p*-value to adopt a more stringent threshold or to adjust the rate of potential false positives. However, there have been concerns regarding correction methods that have been widely implemented in fMRI analysis tools. For instance, research has warned that correction methods implemented in SPM, FSL, and AFNI, clusterwise correction methods in particular, are likely to fail to properly control a false positive rate (Eklund, Nichols, & Knutsson, 2016; Nichols, Eklund, & Knutsson, 2017). Voxelwise inference with correction has been found to be somehow more valid compared with clusterwise inference; however, a recent report demonstrated that voxelwise inference might be conservative and reject true positives (Type II error) (Eklund et al., 2016). Given that assuring a desirable level of statistical power while controlling false positives is also a significant issue in fMRI research (Lieberman & Cunningham, 2009), we intended to address the aforementioned issues in the present study.

We may consider Bayesian statistics as an alternative approach to address the aforementioned issues associated with *p*-values in fMRI research. *P*-values have been used in hypothesis testing within the framework of commonly-employed frequentist statistics (Kyriacou, 2016). As Bayesian statistics is based on statistical assumptions and mechanisms that are fundamentally different from those in frequentist perspective, it is not necessary to use *p*-values in Bayesian hypothesis testing (Gelman & Shalizi, 2013). Thus, Bayesian statistics would be a possible way to avoid issues associated with use (or misuse) of *p*-values in statistical inference (Han & Park, 2018). Unlike frequentists, Bayesians assume that parameters of interest are randomly distributed. Instead of estimating fixed parameters, Bayesians present parameters of interest, such as effect sizes, in terms of probability distributions (Woolrich, 2012). While estimating a specific parameter of interest, Bayesians start with a prior distribution, *P* (*H*), that indicates the hypothetical probability distribution of the parameter. The distribution is being continuously updated through observations. Once the iterative observation process is completed, the parameter of interest is estimated based on the finally updated probability distribution, a posterior distribution, *P* (*H*|*D*). The update process is performed base on Bayes Theorem as follows:

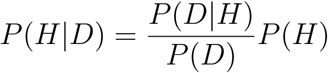

when *H* indicates a hypothesis and D data. In addition, *P* (*D*|*H*) indicates the likelihood that is the probability to observe the data given that *H*, the hypothesis, is true. *P* (*D*) is a probability of data that normalizes the constant of the numerator (Rouder, Speckman, Sun, Morey, & Iverson, 2009).

Following the aforementioned mechanism of Bayesian updating, we can examine the posterior probability of distribution of our hypothesis of interest. For example, if we intend to examine the posterior probability of a null hypothesis, *P* (*H*_0_|*D*), we start with its prior distribution, *P* (*H*_0_) and then update the distribution with collected data. Likewise, we can also examine the posterior probability of our alternative hypothesis, *P* (*H*_1_|*D*), as well (Han & Park, 2018).

With the aforementioned method, we can conduct Bayesian inference to test our hypothesis of interest. For instance, if we intend to examine whether a null hypothesis or alternative hypothesis is more likely to be supported by data, we can compare their posterior probability distributions, which have been updated with collected data. If we have *P* (*H*_0_|*D*) and *P* (*H*_1_|*D*), the posterior probability distribution of a null and alternative hypothesis, respectively, then we can calculate the posterior odds, which represents the ratio of the posterior probability of an alternative hypothesis to that of a null hypothesis as follows:

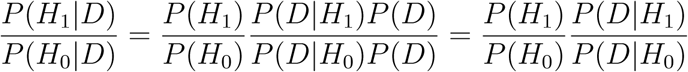

From this equation to estimate the posterior odds, we can extract the Bayes Factor (BF) that has been widely used in Bayesian inference. The BF is the second term in the right-hand side, and it represents the ratio of the amount (or strength) of evidence that is provided by data for either *H*_1_ or *H*_0_ (Rouder et al., 2009). If we are interested in whether and how strongly collected data supports in favor of our alternative hypothesis, then we can examine 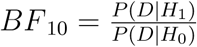. Similarly, if we intend to test whether evidence from data supports in favor of a null hypothesis, we can examine 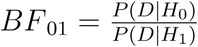. We can determine the strength of supporting evidence and whether our hypothesis of interest is supported by the evidence by examining BFs. There have been several guidelines to interpret BFs in Bayesian inference and hypothesis testing that have been proposed by Bayesian statisticians (see Table S1 for further details).

For example, if we get *BF* _10_ = 15, then we may conclude that evidence strongly supports in favor of *H*_1_ (compared with *H*_0_) and are likely to accept an alternative hypothesis. On the other hand, if a calculated 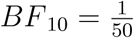, then evidence very strongly supports in favor of *H*_0_ (compared with *H*_1_), and it is plausible to accept a null hypothesis instead of an alternative hypothesis. Unlike *p*-values that have been used in frequentist hypothesis testing and indicate the extremity of data, but do not provide any direct information regarding whether we need to accept or reject a hypothesis, an alternative hypothesis that we are mainly interested in in particular, BFs more directly show us whether or not collected data supports the hypothesis. Given this point, Bayesian inference based on BFs would allow us to more directly test our hypothesis, which could not be easily done within the frequentist framework. Moreover, Bayesian statisticians have argued that it is relatively freer from issues associated with inflated false positives and multiple comparison correction. According to their arguments, it is because Bayesian statistics do not rely on frequentist assumptions associated with false positives but focuses on updating the posterior probability distributions of parameters of interest based on data (Gelman, Hill, & Yajima, 2012; Han & Park, 2018).

Furthermore, recent research has shown that the use of Bayesian methods in fMRI analysis can address the issues associated with inflated positives while maintaining good sensitivity. For instance, Han and Park (2018) examined sensitivity and selectivity of Bayesian and classical second-level fMRI analysis by reanalyzing real fMRI image datasets. Their performance assessment reported that Bayesian analysis can show relatively better sensitivity and selectivity compared with classical analysis. Moreover, a recent study that developed and tested Bayesian fMRI image meta analysis reported that results of Bayesian meta-analysis were more consistent with the results of activation foci syntheses based on large-scale fMRI database compared with classical meta-analysis (Han & Park, 2019). Given these results, we may consider that Bayesian inference in fMRI analysis can be a possible way to deal with the current methodological issues in fMRI research. In fact, Bayesian method has been implemented in fMRI analysis tools, such as SPM (Penny & Friston, 2007), NPBayes-fMRI (Kook, Guindani, Zhang, & Vannucci, 2017), etc.; options to conduct Bayesian analysis are available to end users, fMRI researchers, thanks to presence of the aforementioned tools.

However, the aforementioned Bayesian fMRI analysis studies did not directly control false positive rates although multiple comparison correction has been one of the fundamental issues in the field. Instead, they employed stringent Bayes Factor thresholds to make their analysis more conservative and selective (Han & Park, 2018, 2019). In addition, likewise, the majority of tools that enable researchers to conduct Bayesian analysis do not provide options to perform multiple comparison correction. In fact, Bayesian statisticians have not seriously concerned the issue of multiple comparison correction in Bayesian inference given that Bayesian inference is based on different assumptions compared with classical inference that strongly relies on Type I error rate, *α*. However, there have been debates about whether Bayesian inference should also attempt to directly address the multiple comparison correction issue (Berry & Hochberg, 1999). A group of neuroimaging researchers developed a tool to perform false-discovery rate correction with Bayesian inference (Kook et al., 2017). This tool adjusted posterior probabilities of inclusion to perform the correction. However, they did not assesses the performance of their correction method. In addition, their tool did not support hypothesis testing based on BF. Although some other Bayesian statistics used in the previously developed tools, such as posterior probabilities, may also provide users with information regarding presence of an effect, they would not be suitable to compare the comparative likelihood of *H*_0_ versus that of *H*_1_ that could be done more intuitively with BF (Han & Park, 2019).

To address the aforementioned limitations of prior research regarding Bayesian multiple comparison correction, in the present study, we developed and tested Bayesian multiple comparison correction for second-level fMRI analysis, specifically voxelwise inference. To develop the correction method, we referred to a recent study done by de Jong (2019) that implemented Westfall, Johnson, and Utts (1997)’s Bayesian correction method. de Jong (2019)’s method has been applied in a widely-used Bayesian statistical tool, Jeffreys’s Amazing Statistics Program (JASP) (Love et al., 2017), to perform multiple comparison correction in post-hoc testing. We composed an R script that applied the aforementioned correction method within the context of second-level fMRI analysis, the analysis of contrast images created from first-level analyses. In particular, we intended to focus on the one-sample *t*-test that is the simplest form of *t*-test in order to preliminarily examine whether the correction method could be implemented in fMRI analysis. In addition, to examine whether our method was reliable and credible, we conducted a performance assessment of our method; we particularly focused on selectivity and sensitivity by utilizing false alarm and hit rate as proxies for the aforementioned qualities.

## 2 Method

### 2.1 Materials

We used contrast images created from the first-level (or subject-level) analysis as inputs for our second-level (group-level) analysis. Each input contrast image contained results from the whole-brain tests within each subject. In the case of our study, each voxel in the input contrast image represented the difference in brain activity between two task conditions in that specific voxel. For instance, if brain activities in two conditions, A and B, were compared at the first level, then the value in voxel X in the resultant contrast image of subject Y demonstrated “A-B” in voxel X within subject Y. If a total of *N* participants were examined, then *N* contrast images were used as inputs in our second-level analysis.

We used two different types of datasets, including simulation and real image datasets, to test our Bayesian multiple comparison correction method. First, we created a simulation dataset. The simulation dataset was created by drawing 5mm-radius spheres in an MNI space. The coordinates of the sphere centers were determined by referring to activation foci information reported in Han (2017) (see Table S2 in Han (2017) for coordinates information). All activated voxels in the spheres were marked with 1 while the other background voxels were marked with 0 (see Figure 1 for the created original simulation image). We compared 0 versus 1 in the present study because we intended to assume a true signal (or activation) in the designated voxels. Then, for the evaluation purpose, we added noise to the images based on the created original simulation image by employing the image generation method used in similar prior studies (Han & Park, 2018; Woo, Krishnan, & Wager, 2014). The noise-added images were created by adding Gaussian noise to each voxel in the original simulation image. The standard deviation of the Gaussian noise was adjusted to 50% of the original signal strength (.5), so the Gaussian noise was generated by following *N* (0, .5). To examine whether a sample size influenced analysis results, we prepared noise-added simulation images for five different sample sizes, *N* = 4, 8, 12, 16, and 20. For instance, in the case of *N* = 4, we created four random-noise added images based on the original image.

**Figure 1.**
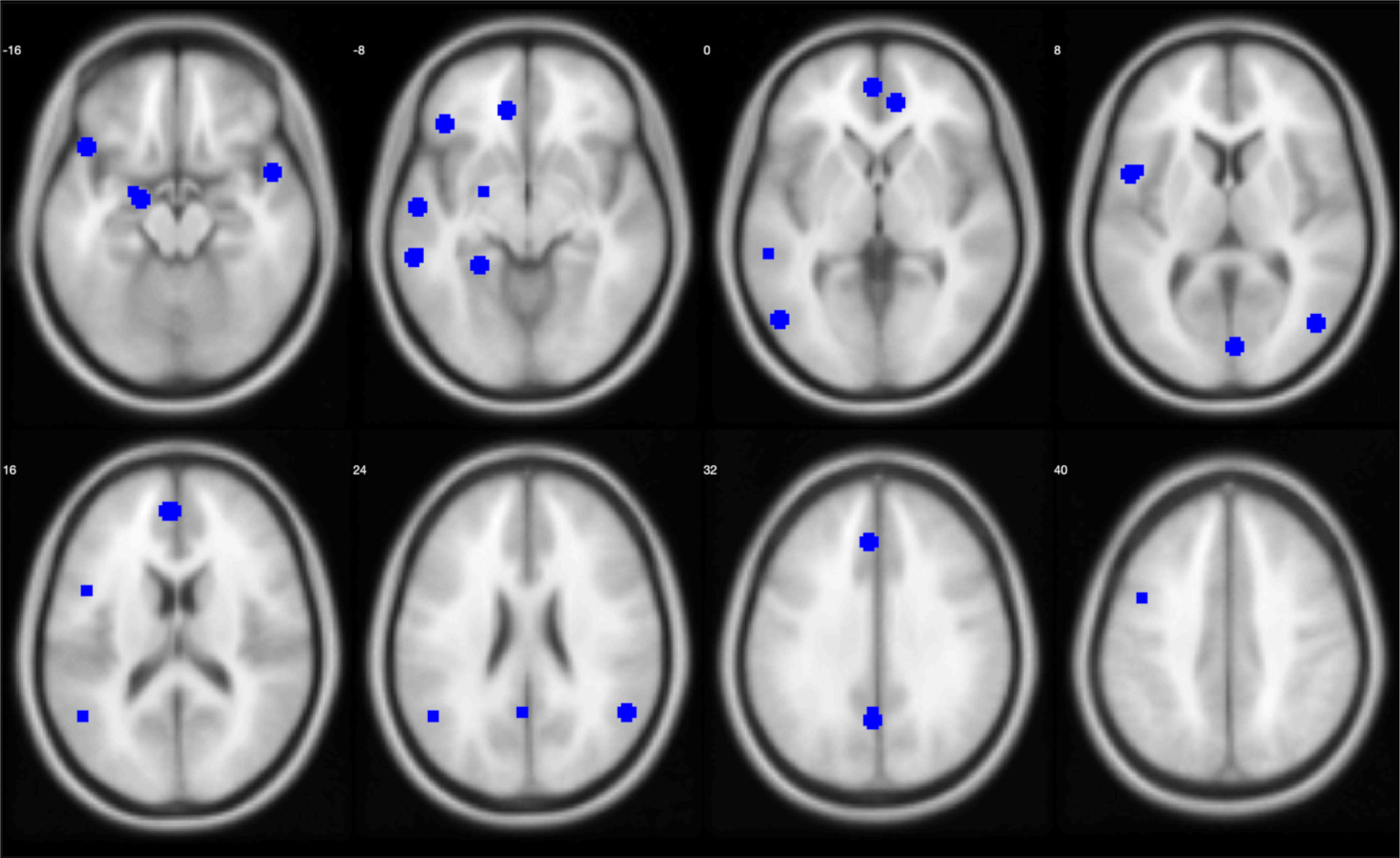
Created original simulation image. Blue voxels (signal strength = 1) represent activated voxels in created spheres. Background voxels were marked with 0.

Second, to examine how the Bayesian multiple comparison correction method worked in the realistic contexts, we prepared real image datasets as well. Five real image datasets were acquired from NeuroVault at https://neurovault.org/ (Gorgolewski et al., 2015) that is a large-scale repository for sharing contrast and statistical fMRI images or lab webpages. We downloaded the aforementioned five datasets that contained contrast images that were created from first-level analysis from the aforementioned sources. Table 1 briefly describes each dataset. The downloaded image files were available in NIfTI format for further analysis. All simulation and real image datasets that were used in the present study are downloadable via Open Science Framework via https://osf.io/dbmu9/.

**Table 1.**
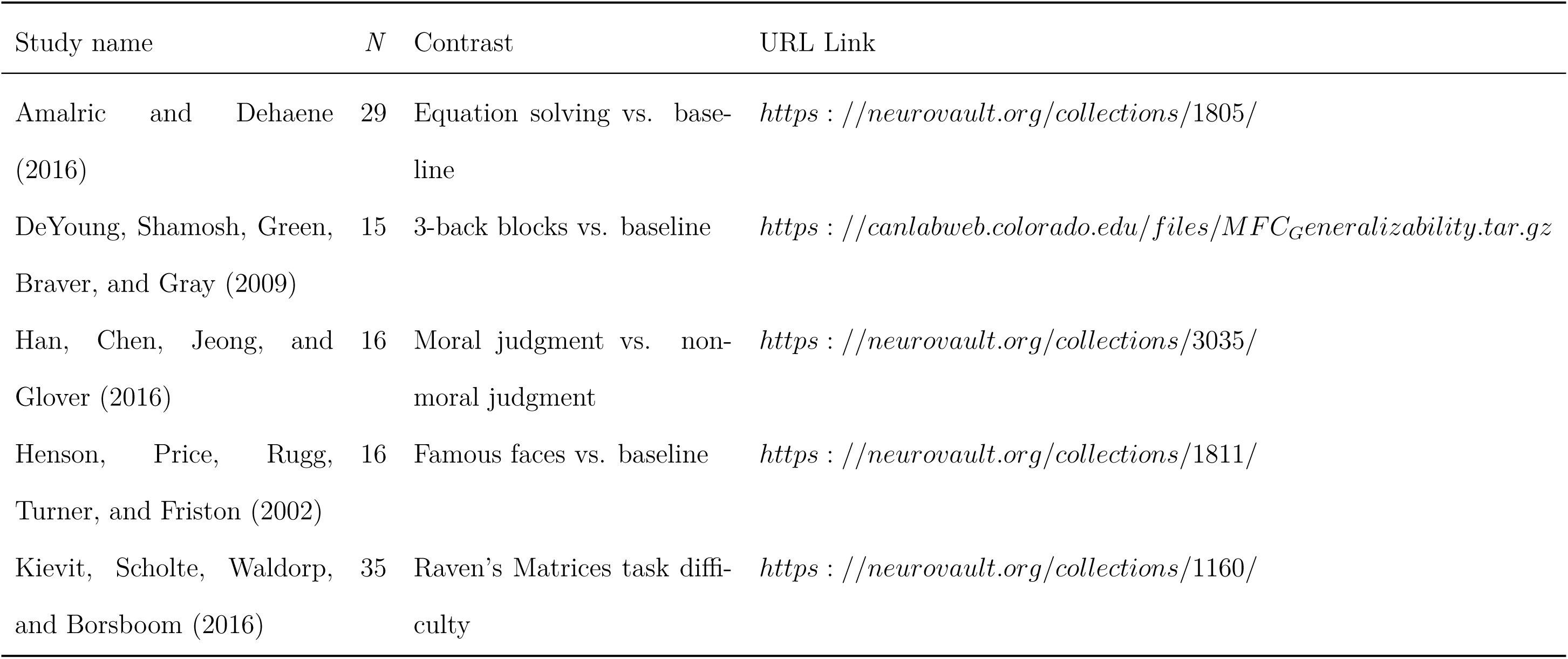
Analyzed real image datasets downloaded from NeuroVault.

### 2.2 Multiple Comparison Correction for Bayesian *t*-test

In the present study, we used contrast images created from the first-level (or subject-level) analysis as inputs. Each input image contained differences in brain activity between different conditions in the whole brain voxels. With the input images, we intended to test whether there was a significant non-zero difference between conditions at the group level. As mentioned in the prior subsection, let us assume that we intended to examine the contrast of “A-B (conditions).” Then, the parameter of our interest would become *δ*, the effect size of the difference “A-B” at the group level. Based on this, we shall set alternative and null hypotheses as follows:

*H*_1_: brain activity in condition A is different from that in condition B (A - B ≠ 0; *δ* ≠ 0)

*H*_0_: brain activity in condition A is not different than that in condition B (A - B = 0; *δ* = 0)

We performed multiple comparison correction in Bayesian inference based on the number of voxels to be tested. According to de Jong (2019) that implemented Bayesian multiple comparison correction in diverse statistical tests, Westfall et al. (1997)’s correction method that takes into account the number of groups (*m*) to be examined shows the better performance compared with the correction method based on Bonferroni correction that is overly conservative. Thus, we also used Westfall et al. (1997)’s method that was implemented in de Jong (2019) in our study.

de Jong (2019) assumed that *m* groups conceptually underlie *k* pairwise comparisons to be performed. The most prevalent form of pairwise comparison is a *t*-test that is also performed in the present study at each voxel. If we assume that a comparison that occurs in each voxel (e.g., brain activity in condition A vs. B) is a pairwise comparison and there are a total of *k* voxels to be analyzed in brain images, then it is possible to say that there are a total of *k* pairwise comparisons to be performed. Then, if we apply Westfall et al. (1997)’s and de Jong (2019)’s approach to this case of the analysis of fMRI images with *k* voxels, we can assume that there are *m* groups to be tested that conceptually underlie *k* pairwise comparisons. Because each individual comparison is a pairwise comparison between two conditions, the relationship between the number of pairwise comparisons (*k*) and that of underlying groups (*m*) can be written in the form of combination. Hence, the relationship between the group number, *m*, and the total number of existing pairwise comparisons, *k*, becomes:

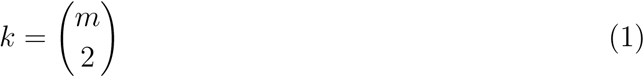

In our study, there were a total of *k* voxels to be examined, so *k* pairwise comparisons, *t*-tests, were performed. Because a *t*-test that compared brain activity in two conditions was performed in each voxel, we assumed that *k* pairwise comparisons were performed in the whole brain. Thus, with (1), we numerically calculated the number of conceptually underlying groups to be tested, *m*, from the *k* number. To search for a positive integer *m* numerically, we attempted to find the smallest *m* that sufficed:

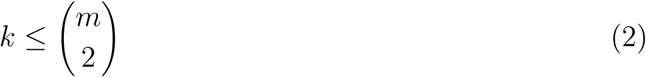

With the numerically calculated *m*, we were able to perform multiple comparison correction by following the method proposed by Westfall et al. (1997) and implemented by de Jong (2019). According to de Jong (2019), the correction of a family-wise error rate can be performed by adjusting the prior distribution in Bayesian inference. Let us assume that the prior distribution of a null hypothesis is *P* (*H*_0_) before correction for pairwise comparison. If we are interested in examining a difference between a global mean, *μ*, and a mean in one of *m* groups to be compared, *μ*_*i*_, in this situation, we can consider two cases related to *μ*_*i*_,

1. *μ*_*i*_ = *μ*, the probability of this case is *τ*, or
2. *μ*_*i*_ ∼ *G, G* is a certain continuous distribution that *μ*_*i*_ from this distribution is never equal to each other, and the probability of this case is 1 − *τ*.

If we are interested in a pairwise comparison between two values, *μ*_*j*_ and *μ*_*k*_, then the probability of whether the two values are identical to each other, *P* (*H*_0*jk*_) should be *τ* × *τ* = *τ* ^2^, because these two values could not be identical to each other as *μ*_*j*_ and *μ*_*k*_ could not be the same in distribution *G* when 2 is the case. If we expand this and consider a hypothetical case when *H*_0_ is true in all *m* underlying groups (e.g., there is no difference in brain activity between conditions A and B in all tested voxels), then *P* (*H*_0_) = P(all *μ*_i_ = *μ*) = *τ* ^*m*^ as de Jong (2019) showed. From this, 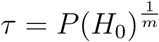. If we intend to get *P* (*H*_0*i*_) that represents the probability that *H*_0_ is true within a specific pairwise comparison, then we can rewrite *P* (*H*_0*i*_) with the aforementioned *P* (*H*_0_) and *τ* as follows:

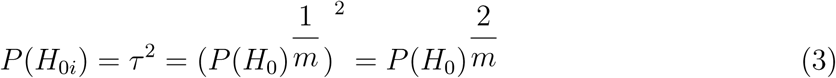

Thus, following (3), for a pairwise comparison, the original prior distribution, *P* (*H*_0_), is adjusted to 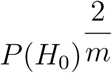. In the present study, for Bayesian multiple comparison correction, we also employed the procedure and used 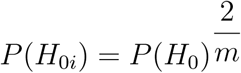 as the adjusted prior distribution for the pairwise *t*-test in each voxel when the original prior distribution was *P* (*H*_0_) and a total of *k* voxels were tested.

For the prior to be used in our Bayesian analysis, we decided to use the Cauchy distribution based on several reasons. First, the Cauchy distribution has been widely used in psychological and neuroscience studies (Gronau et al., 2017); in fact, it has been adopted as the default prior by JASP, a widely used statistical tool for Bayesian inference (Love et al., 2017). Second, the Cauchy prior was used in a prior study that performed Bayesian inference with fMRI images, and the study reported that the performance of Bayesian inference when the Cauchy prior was applied was better than that of classical inference. In addition, the Cauchy prior used in the prior study was robust against the change in the Cauchy prior parameter when the robustness check was conducted. Finally, in terms of survived voxels, when Bayesian thresholding was performed with the Cauchy prior, the pattern of the survived voxels was more similar to the brain activity pattern found from the meta-analysis of large fMRI database compared with when classical inference and thresholding were performed (Han & Park, 2018). Thus, in our study, we used the Cauchy distribution as the prior distribution to conduct second-level analysis.

Following prior studies about Bayesian *t*-test that used the Cauchy distribution that is centered around zero (*x*_0_ = 0) as the prior distribution of a parameter of interest (e.g., the effect size of a difference in brain activity between two conditions, *δ*) (Rouder et al., 2009), let us assume the prior distribution of the null hypothesis as follows:

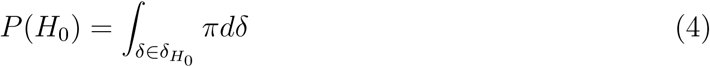

where *π* denotes the probability density on the parameter in the parameter space 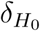 under *H*_0_, and

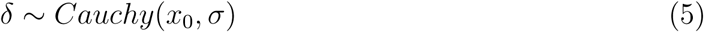

*x*_0_ is the location parameter and *σ* is the scale parameter. In this context, we assume that *x*_0_ = 0 since we are interested in *H*_0_ and a probability distribution that is centered around zero. Prior research suggests that the default value for *σ* is 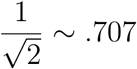 in *t*-tests in the fields of psychological science (Gronau et al., 2017). In fact, this parameter value was used in a previous fMRI analysis that reported the better performance of Bayesian inference compared with classical inference (Han & Park, 2018), so we decided to use this value in our study as well.

Multiple comparison correction and calculate the adjusted prior distribution, *P* (*H*_0*i*_) from *P* (*H*_0_), are done with a predetermined *α* value. Given that *α* represents the probability of rejecting the null hypothesis given that it is true, 1 − *α* would represent the probability of rejecting the null hypothesis given that it is false, the probability of correct rejection. Thus, *α* can be defined as (Weisstein, 2019):

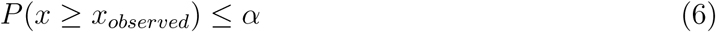

when *x* represents the resultant statistical summary of interest, *δ* in our case, and *x*_*observed*_ what is actually observed. In this case, *P* is conditional on *H*_0_. Accordingly, 1 − *α* should be:

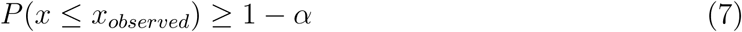

Then, we shall assume that the prior distribution, *P* (*H*_0_), is associated with the probability that an effect does not exist and the distribution of the effect size would be centered at zero (null) (de Jong, 2019), and shall search for at which point of *δ* the calculated *P* (*H*_0_) would become 1 − *α*. From (4) and (6), we need to find *x* that suffices:

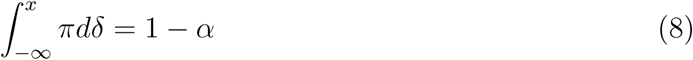

In this context, *x* can be obtained numerically. Once we have *x*, then with (3), find *σ*_*i*_ that suffices:

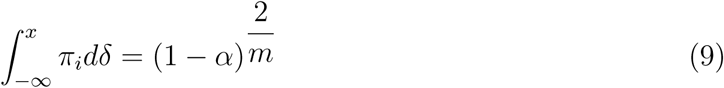

Again, the scale parameter, *σ*_*i*_ can be obtained numerically. Then, in the following *t*-tests, we shall use the adjusted scale parameter, *σ*_*i*_ for the Cauchy prior distribution in order to apply multiple comparison correction.

Here is an example from the analysis of the simulation dataset in the present study. For example, in the case of the aforementioned analysis, the total number of voxels to be tested (*k*) was 91 × 109 × 91 = 902, 629. If *k* = 902, 629, then (2) with the smallest *m* became:

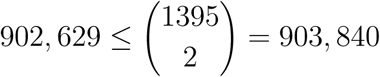

The smallest *m* can be acquired numerically. Thus, *m*, the number of groups that conceptually underlie *k* pairwise tests, was determined to be 1,345 in this case.

Then, we attempted to find the adjusted scale parameter for the Cauchy prior distribution, *σ*_*i*_ with the found *m* value and (8) and (9). Because we intended to use *α* value that has been widely used in the fields of psychology and neuroscience, *α* = .05, with (8), we numerically searched for *x* that sufficed:

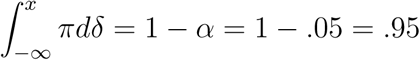

The numerically obtained *x* was 4.46, when 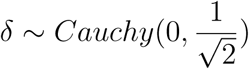, the aforementioned default Cauchy prior, in our simulation. Because we set *α* = .05 and the found *m* was 1,345 in our simulation, from (9),

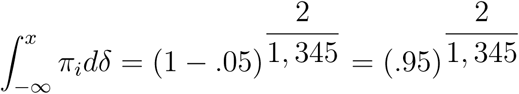

Then, we numerically obtained *σ*_*i*_ = .001, when *δ* ∼ *Cauchy*(0, *σ*_*i*_). Consequently, in the analysis of the simulation dataset, we used *Cauchy*(0, .001) for the prior distribution in our *t*-tests to implement Bayesian multiple comparison correction. Figure S1 demonstrates both the prior distribution before and after adjustment. The adjusted prior distribution became sharper compared with the original distribution. As shown in Figure S2, the sharpened prior distribution contributes to the reduction of calculated BF, so it would make the detection of effects in voxels more stringent and conservative.

### 2.3 Implementation of the Correction Method with R

We composed an R script to implement the Bayesian multiple comparison correction method explained in the prior section. All related source codes and tested data files are shared via Open Science Framework at https://osf.io/dbmu9/. In addition, in supplementary materials, we added a brief tutorial for our Bayesian multiple comparison correction R script.

Our R script was coded to analyze contrast images that were created from the first-level (individual-level) analysis. For instance, when the moral psychology fMRI data was tested, the R script used sixteen contrast images that were created from the first-level analysis that compared neural activity between moral task versus non-moral task conditions in each individual participant as inputs. In addition, to determine *k* in (1), the R script read a mask image that demonstrated voxels that were tested in classical second-level analysis. With the provided mask image, the R script estimated the minimum *m* in (2) (see lines 50-59). Then, the script numerically estimated *x* that suffices (10) once *α* (e.g., *α* = .05) was provided (see lines 68-150). Based on the provided *α*, estimated *x*, and *m*, our R script numerically estimated the adjusted Cauchy scale parameter, *σ*_*i*_, that sufficed (11). Basically, our R script calculates *σ*_*i*_ from the default Cauchy prior scale parameter, 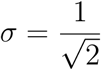. However, when a user specifies the form of the Cauchy prior with a specific *σ* according to the user’s subjective prior belief, then *σ*_*i*_ is calculated from the provided *σ* (see supplementary materials for how to specify *σ* input). Finally, the R script calculated the adjusted Cauchy prior with the estimated *σ*_*i*_.

Then, our R script performed *t*-test in each individual voxel that was specified in the provided mask image. For each voxel, the script calculated a BF to examine whether the provided data supported presence of significant effect. The *t*-test was implemented by an R package named BayesFactor (Morey, Rouder, Jamil, Urbanek, & Ly, 2018). Our R script utilized a functionality implemented in this package, ttestBF, to perform *t*-tests. Although ttestBf implements various types of *t*-tests including one-sample, paired-sample, and independent-sample *t*-tests, we used one-sample *t*-tests in the present study. Contrast values in a specific voxel that were extracted from provided first-level contrast images files were used as inputs to ttestBF. Moreover, the adjusted Cauchy distribution, *Cauchy*(*δ*; 0, *σ*_*i*_), was used as the prior distribution in Bayesian *t*-tests. As a result, the median value of posterior effect size, *δ*, which was a parameter of interest in our study, and BF were calculated for each voxel. Once the median posterior effect size and BF values were calculated for all voxels specified in the mask image, our R script created two NIfTI files recording resultant median posterior effect size and BF values from the performed second-level analysis. We performed the aforementioned Bayesian correction and analysis on UAHPC, a research computing cluster at the University of Alabama, with better computational power.

Bayesian thresholding was performed by assessing whether a calculated BF value in a specific voxel exceeded a pre-determined BF threshold to determine whether the voxel was significantly activated. Following the guideline that was provided in Kass and Raftery (1995) and was used in previous studies that implemented the similar *t*-test method (Han & Park, 2018; Han, Park, & Thoma, 2018; Wagenmakers et al., 2018), we extracted voxels that reported BF greater than 3. According to the guideline and prior research, *BF* ≥ 3 indicates presence of evidence that positively supports a significant effect (or difference). We created thresholded images by using the aforementioned BF threshold with XjView (Cui, Li, & Song, 2015).

### 2.4 Performance Evaluation

We examined the performance of the proposed Bayesian multiple comparison correction by conducting second-level analysis with the simulation and real image datasets that were explained in the materials section. First, we examined the overall performance of the Bayesian multiple comparison method with the simulation dataset. As indicators for performance evaluation, we used two indicators, false alarm and hit rates. These indicators were initially developed to assess the performances of fMRI analysis methods, including classical parametric, non-parametric, and Bayesian analysis methods, with real image datasets in prior research (Han & Glenn, 2018; Han, Glenn, & Dawson, 2019; Han & Park, 2018). The false alarm rate can be used as an indicator of selectivity and be defined in terms of the ratio of voxels that survive thresholding but in fact are inactive. Given that this indicator is associated with the false positive rate, the lower false alarm rate, the better selectivity and overall performance. On the other hand, the hit rate indicates the overall sensitivity of a specific method. The hit rate can be calculated from the ratio of voxels that survive thresholding and in fact are actually active. Thus, the higher hit rate indicates a better sensitivity and overall performance.

Given that the original simulation image file consists of 0 and 1 that indicate absence and presence of activity, respectively, we were able to calculate false alarm and hit rates by comparing the original image and the thresholded image that demonstrated the result of second-level analysis that was conducted with noise-added images. To quantify the aforementioned false alarm and hit rates, we compared the original simulation image with the thresholded resultant images. For the assessment of the performance of our Bayesian multiple comparison correction method, we compared the original simulation image with the thresholded images that demonstrated voxels with *BF* ≥ 3. Then, we also calculated the performance indicators for the classical inference method as well. We compared the original simulation image with the thresholded images that survived voxelwise thresholding (with FWE correction) implemented in SPM 12 at *p* < .05. Furthermore, we also calculated false alarm and hit rates when the multiple correction was not performed in both the cases of Bayesian and classical inference. We repeated the aforementioned evaluation process ten times per case. We used five different sample sizes, *N* = 4, 8, 12, 16, and 20. As an auxiliary test, we examined how BFs changed when the Cauchy prior scale parameter, *σ*, changed as our R script allows a user to enter a user-specified *σ*. Following the prior robustness check in Han and Park (2019), we examined the change in BFs when *σ* changed from .025 to 1.5. This test was done in two specific voxels, one was active (1) and the other was not active (0), in the provided tutorial image data. The true active voxel was located at (45, 89, 36) and the inactive voxel was at (46, 64, 37). We were mainly interested in whether Bayesian *t*-test after applying our multiple correction method produced significantly changed BFs as we altered the *σ* input.

Second, we performed the Bayesian *t*-test with multiple comparison correction with the five real image datasets. We conducted the additional analysis with the real datasets to examine whether our method can also be applicable to different datasets with different image sizes, signal strengths, and noise strengths in more realistic settings. Similar to the case of the examination of the simulation dataset, we analyzed downloaded first-level contrast images with our R script. In addition, we conducted classical *t*-test by using voxelwise inference with FWE correction implemented in SPM 12. Then, we compared the number of voxels survived thresholding in each case. We also examined the number of survived voxels when Bayesian and classical inference were conducted without multiple comparison correction.

## 3 Results

### 3.1 Simulation Image Data Analysis

Figure 2 demonstrates the resultant false alarm and hit rates from the analyses of the simulation image dataset. In addition, Figure 3 shows voxels survived each thresholding method across different sample sizes. The false alarm and hit rates are demonstrated for four different thresholding methods and five different sample sizes. In all cases, from *N* = 4 to 20, Bayesian inference with multiple comparison correction reported the lower false alarm rate compared with both Bayesian and classical inference without correction, but the higher false alarm rate compared with classical voxelwise inference with FWE correction. The hit rate was higher in general when correction was not performed; however, when N ≥ 16, the hit rate became similar or even higher when Bayesian inference was performed with correction compared with when Bayesian inference was performed without correction. When multiple comparison correction was performed, Bayesian inference showed the higher hit rate compared with classical inference. The aforementioned trend of hit rate was consistent across different sample sizes from *N* = 4 to 20.

**Figure 2.**
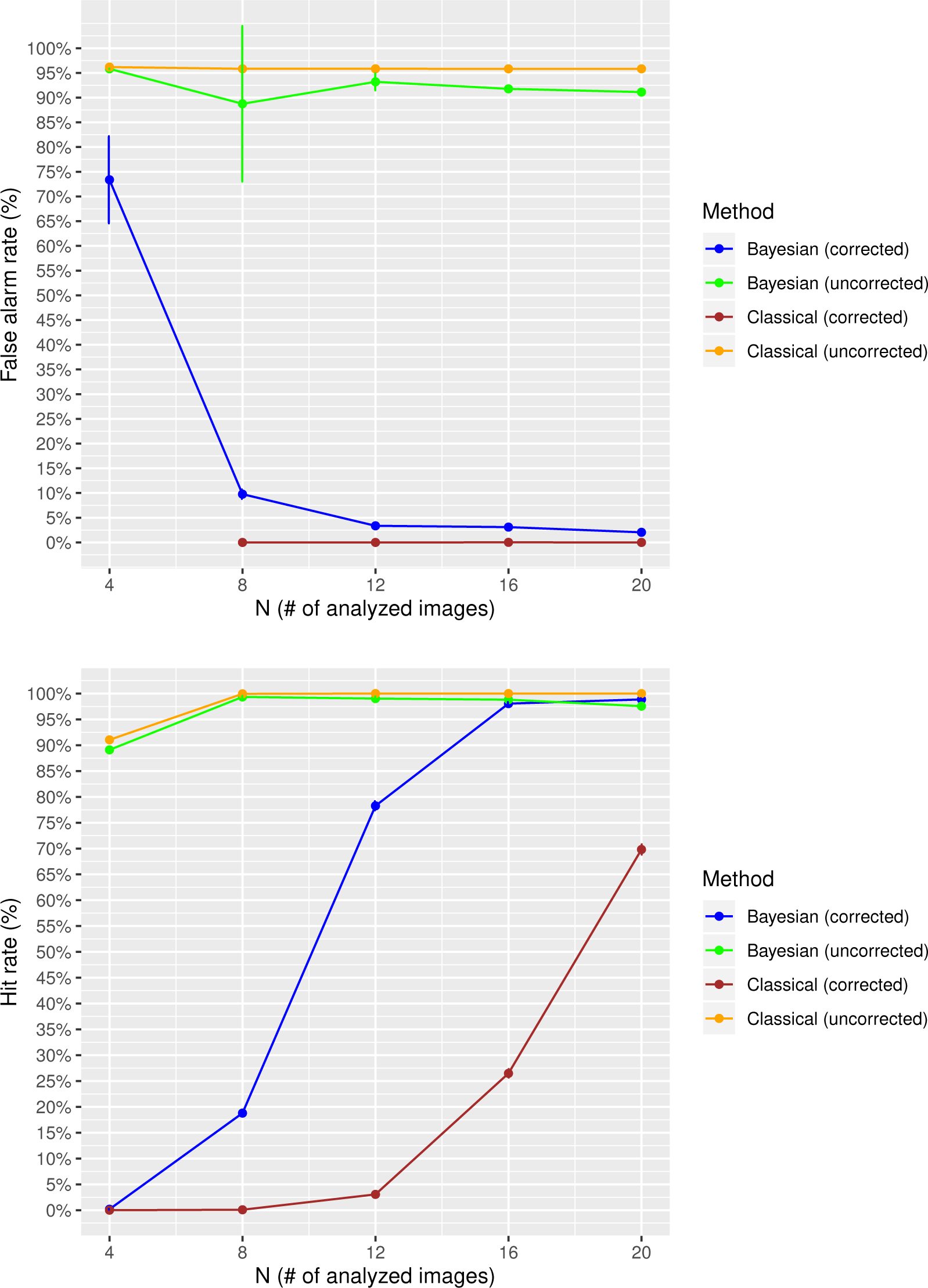
False alarm (top) and hit rates (bottom) in different *N*s. *Note.* False alarm rate of classical inference with correction was omitted at *N* = 4 since no voxel survived with this sample size.

**Figure 3.**
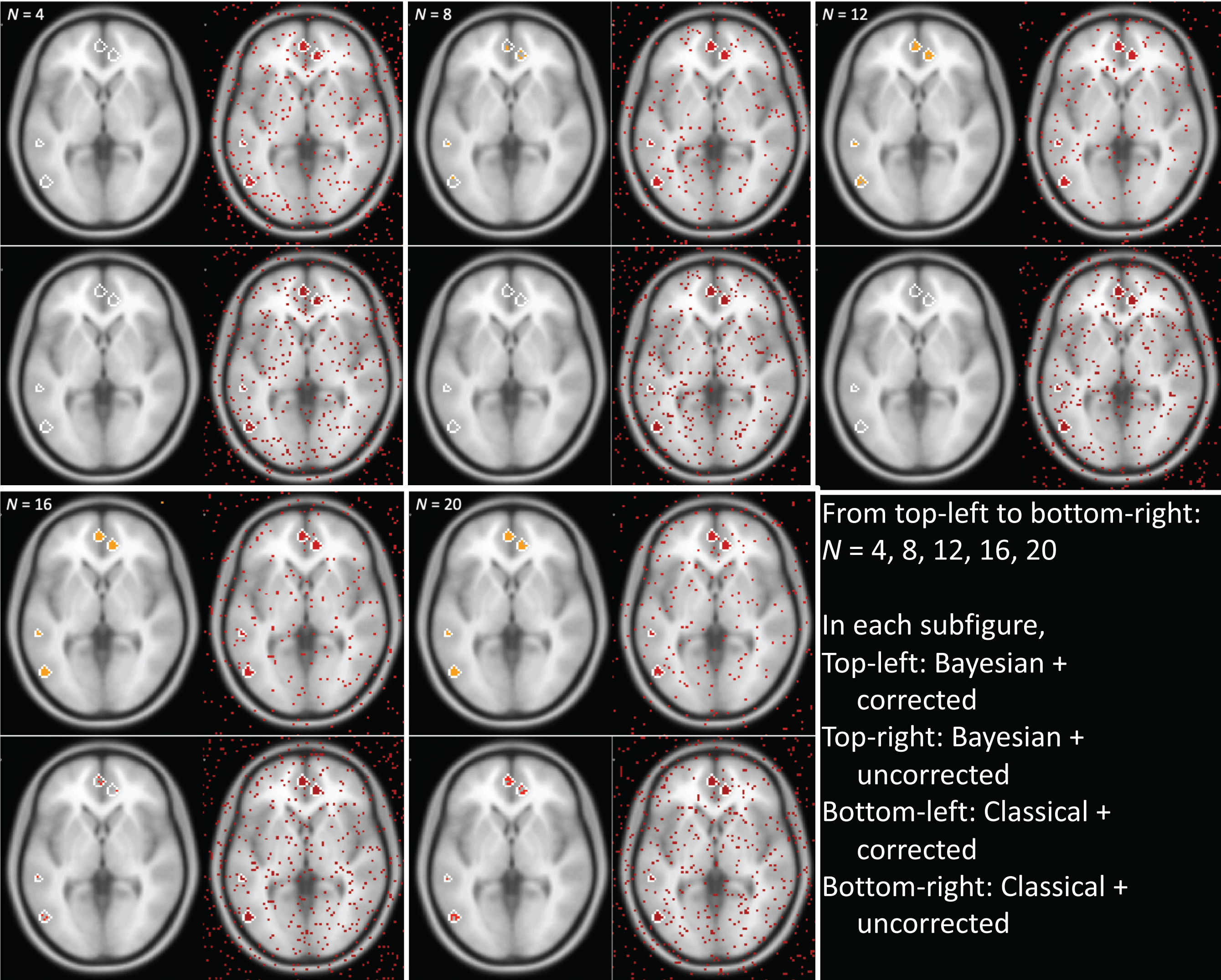
Voxels survived each thresholding method across different sample sizes. White lines: regions contain true positives in the simulated image. Color dots: voxels survived each thresholding method.

The auxiliary test reported that the resultant BF did not significantly change even with the change in the *σ* input. As shown in Figure S3, first, when the tested voxel was originally active, the resultant BF was greater than 3, the BF threshold used in the present study, regardless of the change in *σ* (top figure). Second, when the inactive voxel was tested, regardless of the change in *σ*, the resultant BFs were always smaller than 3 (bottom figure). Hence, in the case of the analysis of our simulation image, the resultant BF did not significantly change even with the change in *σ* that was associated with the change in a subjective prior belief.

### 3.2 Real Image Data Analysis

Results from the analyses of five real image datasets are reported in Table 2. Table 2 demonstrates the number of tested voxels (*k* in (1)) and the number of survived voxels when each thresholding method was applied. In addition, images that demonstrate thresholding result are presented in Figures S4-S8 in supplementary materials. When Bayesian inference with multiple comparison correction was conducted, the number of survived voxels was greater compared when classical voxelwise inference with FWE correction was conducted; the number was smaller compared with when multiple comparison correction was not applied. The aforementioned trend was found from analyses of all five real image datasets.

**Table 2.** Number of tested voxels and number of survived voxels from analyses of real image datasets.

| Study name | Voxels<br>tested | Bayesian<br>(corrected) | Bayesian<br>(uncorr.) | Classical<br>(corrected) | Classical<br>(uncorr.) |
| --- | --- | --- | --- | --- | --- |
| Amalric and Dehaene<br>(2016) | 51,360 | 11,664 | 22,608 | 5,016 | 27,256 |
| DeYoung, Shamosh, Green,<br>Braver, and Gray (2009) | 206,777 | 5,524 | 39,636 | 89 | 47,428 |
| Han, Chen, Jeong, and<br>Glover (2016) | 180,401 | 5,491 | 44,098 | 264 | 55,928 |
| Henson, Price, Rugg,<br>Turner, and Friston (2002) | 51,199 | 4,551 | 13,551 | 1,085 | 15,872 |
| Kievit, Scholte, Waldorp,<br>and Borsboom (2016) | 326,293 | 345 | 6,449 | 30 | 12,360 |

## 4 Discussion

In the present study, we implemented Bayesian multiple comparison correction for second-level fMRI analysis based on Westfall et al. (1997)’s correction method implemented by de Jong (2019) in our customized R script. To examine how the developed Bayesian multiple comparison correction worked with fMRI datasets, we tested the method with simulation and real image datasets. To assess the performance of the method, we calculated false alarm and hit rates that are associated with selectivity and sensitivity, respectively, with the simulation dataset. We compared the performances of analysis methods in different conditions, Bayesian versus classical inference and with versus without correction by using the aforementioned two indicators. In addition, we compared results from analyses of the five real image datasets among different analysis methods.

When we conducted Bayesian second-level inference with multiple comparison correction with the simulation dataset, the result suggested that Bayesian inference with multiple comparison correction showed good sensitivity given the resultant hit rate. Our Bayesian method outperformed the classical voxelwise FWE correction method in terms of the sensitivity, which was quantified in terms of hit rate, particularly when the sample size was small (hit rate > 75%, when *N* ≥ 12). The classical voxelwise FWE correction demonstrated the relatively worse hit rate; even when *N* = 20, the resultant hit rate was lower than 70%. Meanwhile, although the classical voxelwise FWE correction showed nearly perfect sensitivity, false alarm rate ∼ 0%, our Bayesian correction method was also able to show good sensitivity, alarm rate < 10%, when *N* ≥ 8. Given these reported results, we shall conclude that our Bayesian correction method was able to produce the better sensitivity compared with the classical voxelwise FWE correction method while maintaining the reasonable sensitivity when the sample size was relatively small. Although the classical method was able to perfectly reject false positives, it was more likely to reject true positives compared with our Bayesian method and could not show good sensitivity. Thus, Bayesian inference with correction can be a reasonable alternative way for second-level fMRI analysis to maximize sensitivity while maintaining the acceptable level of selectivity when a sample size is small.

The similar trend was found in the analyses of real image datasets. In all cases, the number of survived voxels were greater when our Bayesian multiple comparison correction was performed compared with when classical voxelwise FWE correction was performed. Such a trend was consistent even the image size, which was related with the number of comparisons and tests, varied across different datasets. This trend suggests that our Bayesian method is more sensitive compared with the classical method in detecting active voxels. Still, relatively less voxels survived when Bayesian hypothesis testing was conducted with multiple comparison correction compared with when no correction was performed. These results might support the point that Bayesian multiple comparison correction can contribute to controlling possible false positives and maintaining selectivity. In fact, Bayesian methods in general, such as Bayesian structural equation models (Lee & Song, 2004) and Bayesian multi-level model analysis (McNeish & Stapleton, 2016; Stegmueller, 2013), are less likely to produce biased analysis results specifically when a sample size is small compared with classical inference methods given prior research. Hence, the result of the present study, good sensitivity and acceptable selectivity that resulted from use of Bayesian inference with correction when *N* was small, would be consistent with what have been found in the aforementioned previous studies used Bayesian methods.

Furthermore, researchers should also consider epistemological benefits of Bayesian inference as well. Hypothesis testing based on *p*-values could not provide us with any direct information regarding whether our hypothesis (e.g., presence of significant effects) is supported by evidence; instead, *p*-values can only indicate the extremity of data. Instead, calculated BF values directly indicate whether the hypothesis is supported by observed evidence; more specifically, the values (e.g., *BF* _10_) demonstrate whether evidence supports in favor of an alternative hypothesis (*H*_1_) (compared with a null hypothesis (*H*_0_)) (Han & Park, 2018; Rouder et al., 2009). Because one of the major issues associated with use of BFs in neuroimaging research, whether Bayesian methods allow users to perform multiple comparison correction, has been addressed in the present study, Bayesian inference based on BFs can be a useful alternative method in neuroimaging research that possesses the aforementioned epistemological beliefs and can minimize false positives.

In addition, our study made a practical contribution to the field as well. We composed an R script that implemented Bayesian multiple comparison correction in second-level fMRI analysis and made it available to the public via the Open Science Framework (all source codes and tested image data files are available at https://osf.io/dbmu9/). We also provided a brief description regarding how to customize the composed R script so that other researchers conduct their own second-level fMRI data analysis in the repository. Thus, by sharing the customizable source code and tested image datasets that can be used for tutorial purpose, researchers in the field of neuroimaging will also be able to feasibly test and employ the novel analysis method that we developed in the present study.

However, there are several limitations that warrant future studies. First, given that Bayesian inference is performed through iterative Markov Chain Monte Carlo computation processes (Gelman & Rubin, 1992), the implementation of the proposed method requires strong computational power. Given that the aforementioned computational processes should be done in more than hundred thousand individual voxels in fMRI analysis, it took significantly longer time to conduct Bayesian inference compared with classical inference. In fact, we needed to wait about twelve hours until the completion of the computation processes. Because R programs are performed in a single core by default (Gillespie & Lovelace, 2016), it would require long time to complete computationally intensive tasks in general. Thus, it is necessary to consider adopting the multicore programming technique in future studies to address this issue. Second, although we provide researchers with a customizable R script so that they conduct their own fMRI analysis by modifying the provided source code, knowledge in R programming is required to utilize the method proposed in the present study. Hence, it would be necessary to develop a user-friendly interface so that researchers without sufficient programming skills can employ our method while feasibly modifying parameters and options as they intend. Third, in the present study, we only focused on the simplest test, an one-sample *t*-test, since we were mainly interested in the comparison of Bayesian and classical inference with correction in general. In addition, because the one-sample *t*-test at the second level can answer the majority of research questions related to within-subject differences and can be a versatile inference method by using results from diverse types of first-level tests as input contrast image data (Pauli et al., 2016), we decided to focus on this test in our study. Future studies may test different types of statistical tests, such as a two-sample *t*-test, AN(C)OVA, correlation analysis, etc., to examine whether Bayesian correction method performs well in diverse settings. Given that BayesFactor package provides diverse types of Bayesian hypothesis testing, this limitation can be addressed in future studies by customizing the R script. Fourth, we only focused on voxelwise inference in the present study, so we could not consider how to perform clusterwise Bayesian inference with correction in this study. Thus, it would be necessary to consider how to implement Bayesian inference for clusterwise inference in the future research.

## 5 Conclusion

In the present study, we implemented and tested Bayesian multiple comparison correction in second-level fMRI analysis, voxelwise inference in particular, with R. The developed R code and tested data files are shared via the Open Science Framework so that neuroimaging researchers conduct Bayesian second-level analysis with correction with their own dataset through the customization of the provided script. Our Bayesian correction method reported good sensitivity and acceptable selectivity as shown in the resultant hit and false alarm rates particularly when the sample size was relatively small (*N* ≤ 20). Given these, our method will be able to contribute to the field by providing an enhanced alternative multiple comparison method for second-level fMRI analysis.

## Supplementary Materials

### A Brief Tutorial for Bayesian Multiple Comparison Correction R Script

Readers can learn how to use Bayesian multiple comparison correction with the provided R script file and tutorial image data files. For this tutorial, readers need to download all required files from our OSF project page. In the page, https://osf.io/dbmu9/, all tutorial-related files are uploaded in a folder named “***Tutorial***.” Here is a list of files to be downloaded:

***Bayes_test.R → R script file for Bayesian multiple comparison correction***

***Mask.nii → A mask nifti file that specifies which voxels should be tested***

***1.nii - 16.nii → Sixteen contrast image files to be used as inputs***

***List.csv → a list file that specified which contrast images files shall be used as inputs***

The tutorial dataset consists of the simulation data used in the present study. The current version uses a total of sixteen contrast images files as inputs (*N* = 16). In addition, a mask image file (mask.nii) is also required. This mask file can be created by performing classical inference with SPM or other tools. For this tutorial, we uploaded a mask.nii file that was created with SPM 12 with the sixteen image files, 1.nii to 16.nii. This mask file specifies voxels to be tested; voxels to be tested are marked with “1” while those not to be tested are marked with “0” or “NaN.” Here is the 3D view of the currently included mask file, mask.nii:

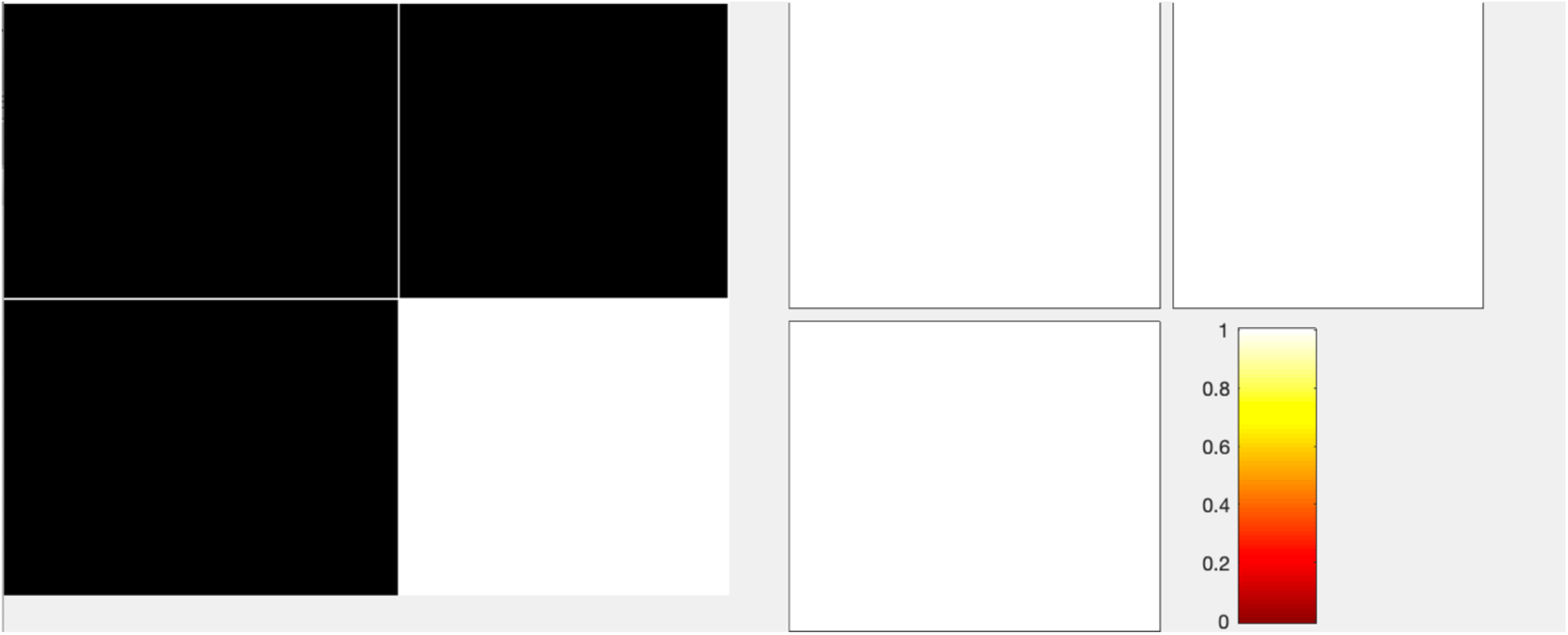

For the readers’ information, here is the 3D view of the first contrast image file to be used (1.nii):

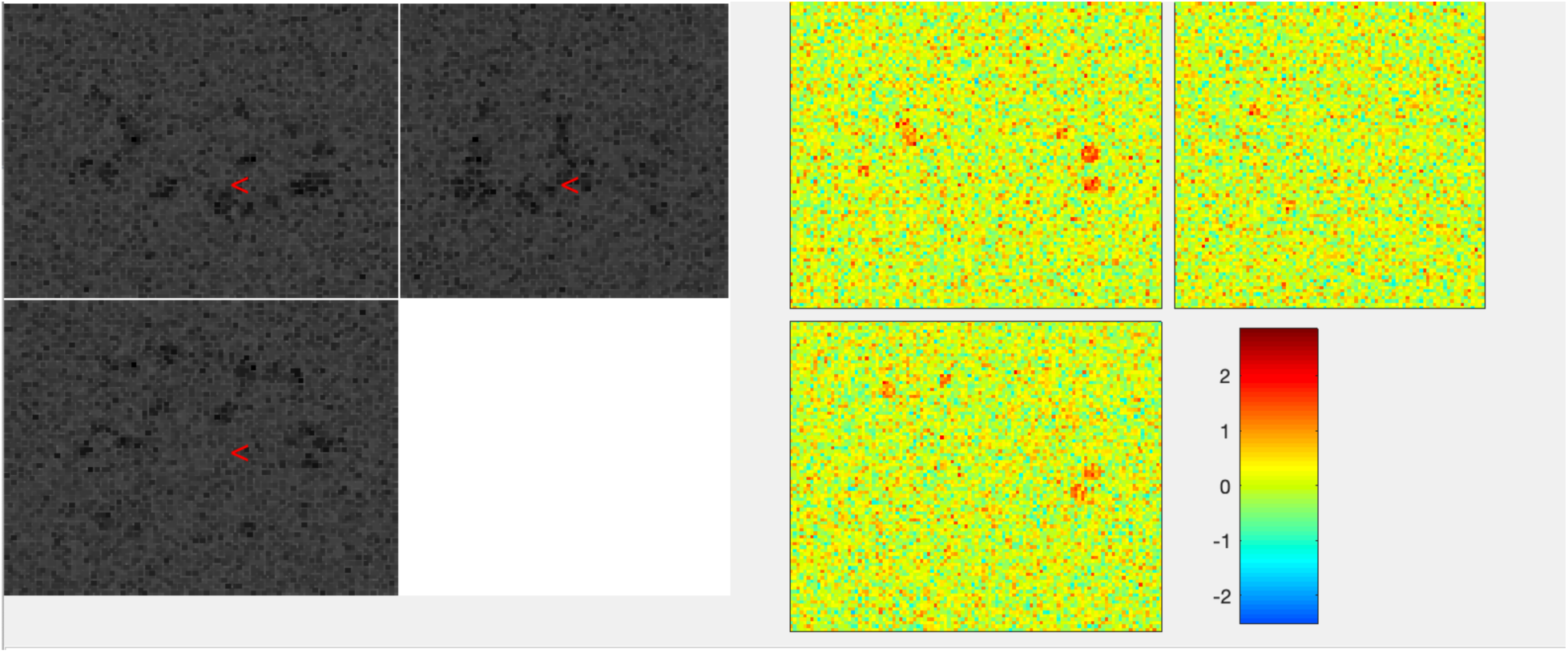

List.csv contains the list of nii files to be analyzed. In the case of this tutorial, it contains filenames 1-16.nii. If a reader intends to analyze his/her own contrast image files, he/she needs to create a new list.csv file accordingly. The content of the csv file is presented below:

Filename

1. nii
2. nii
3. nii
4. nii
5. nii
6. nii
7. nii
8. nii
9. nii
10. nii
11. nii
12. nii
13. nii
14. nii
15. nii
16. nii

The file contains only one column, “***Filename***.” All input contrast image files should be listed in rows below the first row with the column name, “***Filename***.”

In order to run the script, all the downloaded files should be placed in the same folder. R version should be higher than 3. Before run Bayes_test.R, several R packages should be installed. First, in order to run Bayesian inference in R, ***RJAGS*** should be installed. Here is a link to ***RJAGS*** project page for further details: https://cran.r-project.org/web/packages/rjags/index.html. In addition to ***RJAGS***, in R, two packages, “***BayesFactor***” and “***oro.nifti***,” should be installed. These two packages can be installed with “***install.packages***” command in R.

Once all required files are downloaded and required packages are installed, Bayes_test.R can be ran in a command window (Windows) or terminal (MacOS or LINUX). Bayes_test.R requires one optional parameter, whether or not perform Bayesian multiple comparison correction. When no parameter is provided, Bayes_test.R performs correction with the default Cauchy prior scale parameter, 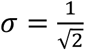 (default option). To run Bayes_test.R with the default option (performing correction), type and run either

***Rscript --vanilla Bayes_test.R***

Or

***Rscript --vanilla Bayes_test.R 1***

In a command line. To run Bayes_test.R without correction with the scale parameter, type and run

***Rscript --vanilla Bayes_test.R 0***

If a reader intends to modify the form of the Cauchy prior with a specific scale parameter based on a subjective belief, the scale can be specified as the third option. For instance, if an intended scale parameter is .5 then type and run either

***Rscript --vanilla Bayes_test.R 1 <u>.5</u>* (**with multiple comparison correction)

Or

***Rscript --vanilla Bayes_test.R 0 <u>.5</u>* (**without multiple comparison correction)

Because Bayesian inference involves iterative processes, it may take up to a couple of hours to complete image analysis; the duration may vary according to the number of voxels to be tested, which is specified in mask.nii, and the performance of a reader’s computer. Once all processes are done, output files are created. If a reader enabled the option to perform correction, two output files, BFs.nii and Ds.nii, are created. BFs.nii contains a resultant Bayes Factor value (after correction) in each voxel and Ds.nii contains a calculated effect size value in each voxel. The 3D view of BFs.nii that will be created from the tutorial input data files looks like:

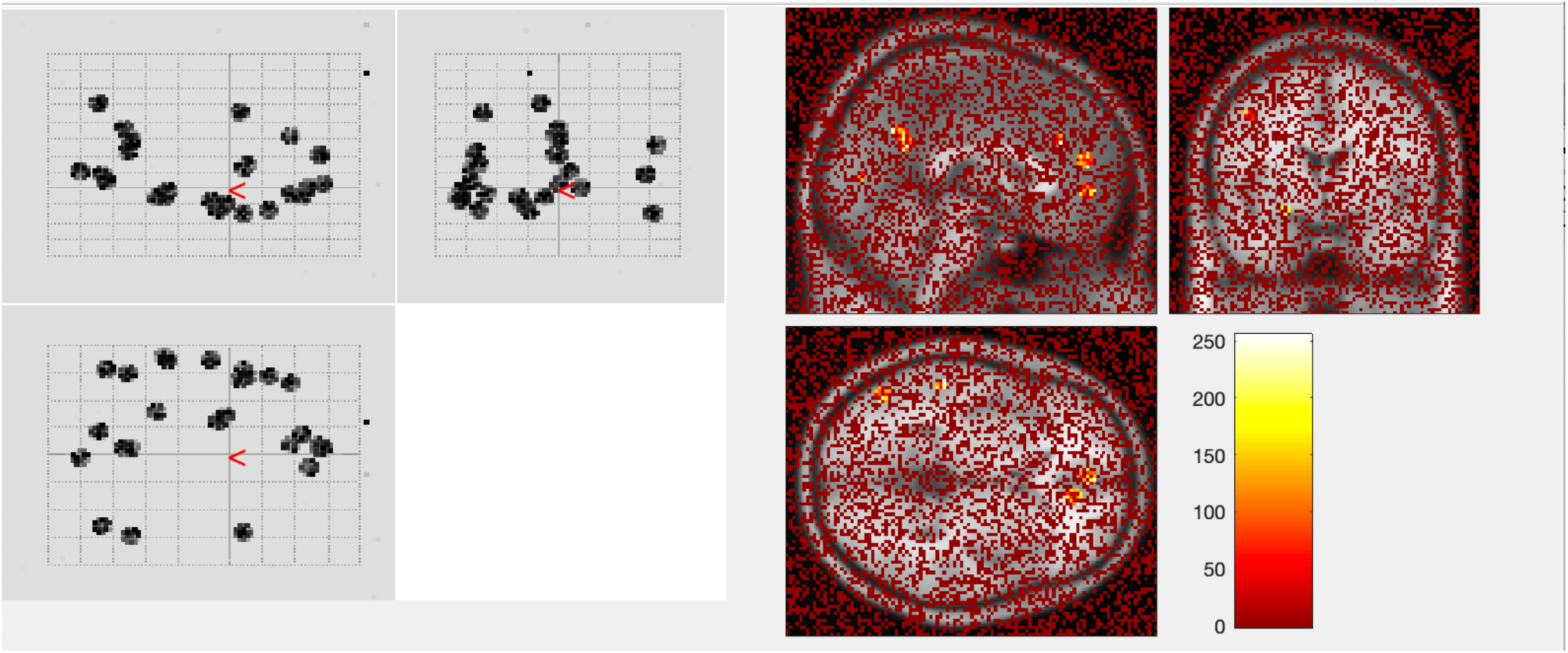

If a reader thresholds this image with a viewer that supports thresholding (e.g., xjview), then he/she will be able to get a thresholded result that is similar to what reported in our paper. If the user applies a Bayes Factor threshold BF ≥ 3, then the resultant thresholded image looks like:

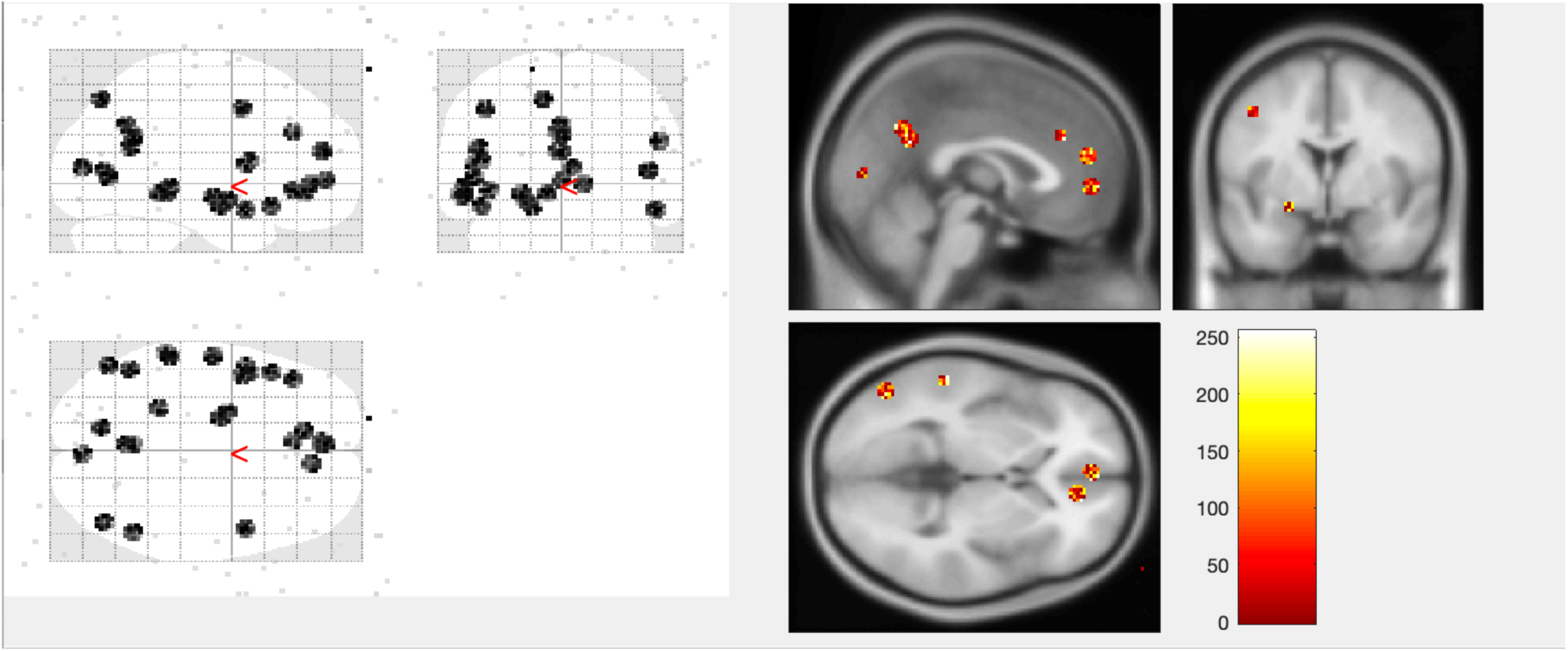

As shown above, when correction is applied and the ordinary Bayes Factor threshold, BF = 3, is applied, our Bayesian multiple comparison correction method was able to well distinguish the true simulated positives as reported in our paper.

If a reader runs Bayes_test.R without correction, then two output files are created. BFs_uncorr.nii contains a calculated Bayes Factor value in each voxel without correction. Ds.uncorr.nii contains a calculated effect size value in each voxel accordingly. BFs_uncorr.nii can also be thresholded with a viewer with a thresholding functionality, such as xjview, similar to the case of BFs.nii explained above.

## Supplementary Tables

**Table S1.** Guidelines to interpret BFs in Bayesian hypothesis testing. Modified from Andraszewicz et al. (2015).

| $BF_{10}$ | | | How to interpret $BF_{10}$ |
| --- | --- | --- | --- |
| | > | 100 | Extremely strong evidence supporting in favor of $H_1$ |
| 30 | to | 100 | Very strong evidence supporting in favor of $H_1$ |
| 10 | to | 30 | Strong evidence supporting in favor of $H_1$ |
| 3 | to | 10 | Moderately positive evidence supporting in favor of $H_1$ |
| 1 | to | 3 | Anecdotal evidence supporting in favor of $H_1$ |
|  | 1 |  | No evidence |
| $\frac{1}{3}$ | to | 1 | Anecdotal evidence supporting in favor of $H_0$ |
| $\frac{1}{10}$ | to | $\frac{1}{3}$ | Moderately positive evidence supporting in favor of $H_0$ |
| $\frac{1}{30}$ | to | $\frac{1}{10}$ | Strong evidence supporting in favor of $H_0$ |
| $\frac{1}{100}$ | to | $\frac{1}{30}$ | Very strong evidence supporting in favor of $H_0$ |
| | < | $\frac{1}{100}$ | Extremely strong evidence supporting in favor of $H_0$ |

## Supplementary Figures

**Figure S1.**
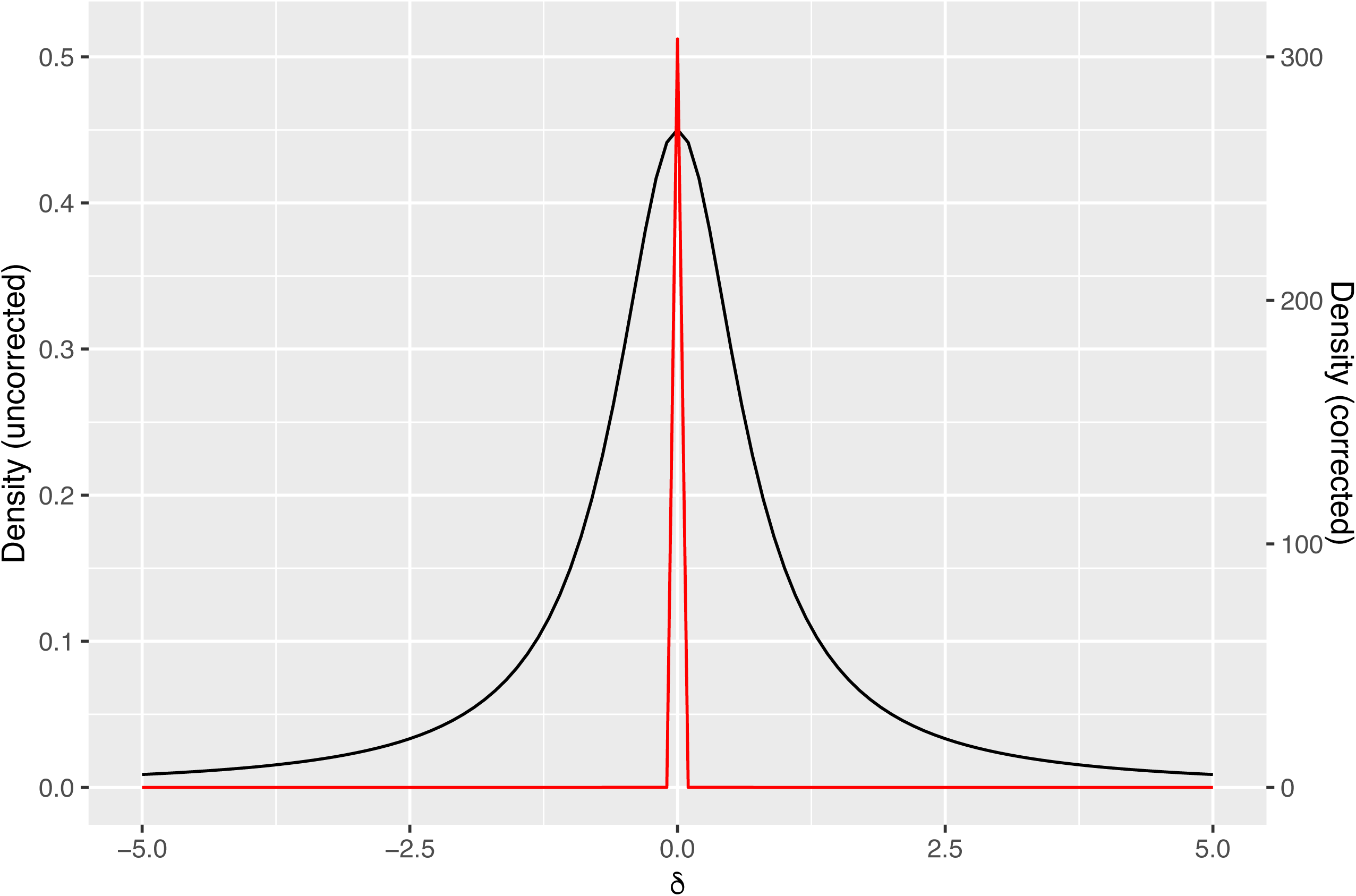
Cauchy prior distributions before and after correction, when m = 1,345. Black: before correction (σ = .707). Red: after correction (σ_i_ = .001).

**Figure S2.**
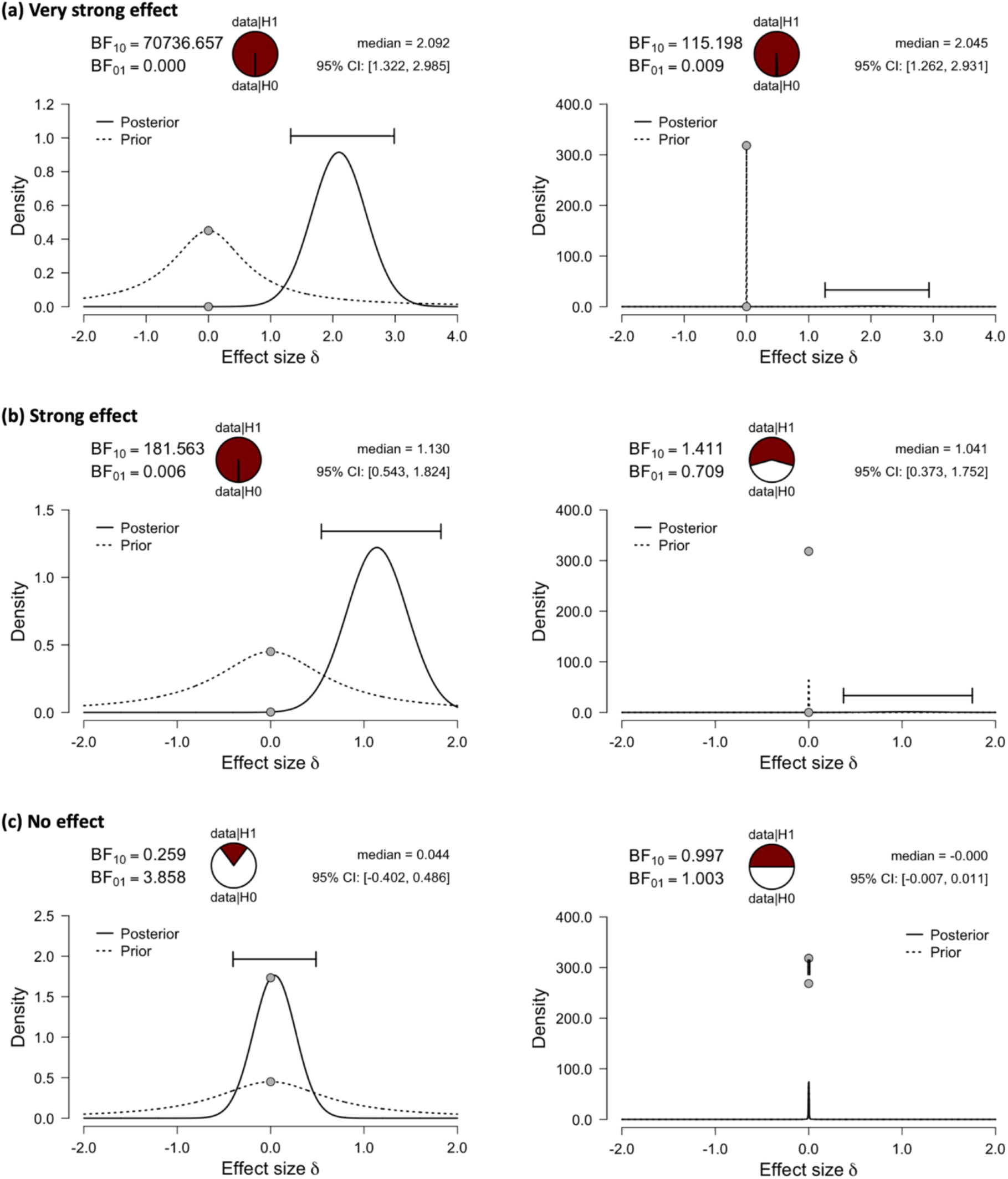
Prior and posterior distributions, and calculated BFs in three different cases when: (a) a very strong effect exists; (b) a strong effect exists; and (c) no effect exists. Left: results without multiple comparison correction. Right: results with multiple comparison correction. Figures created with JASP.

**Figure S3.**
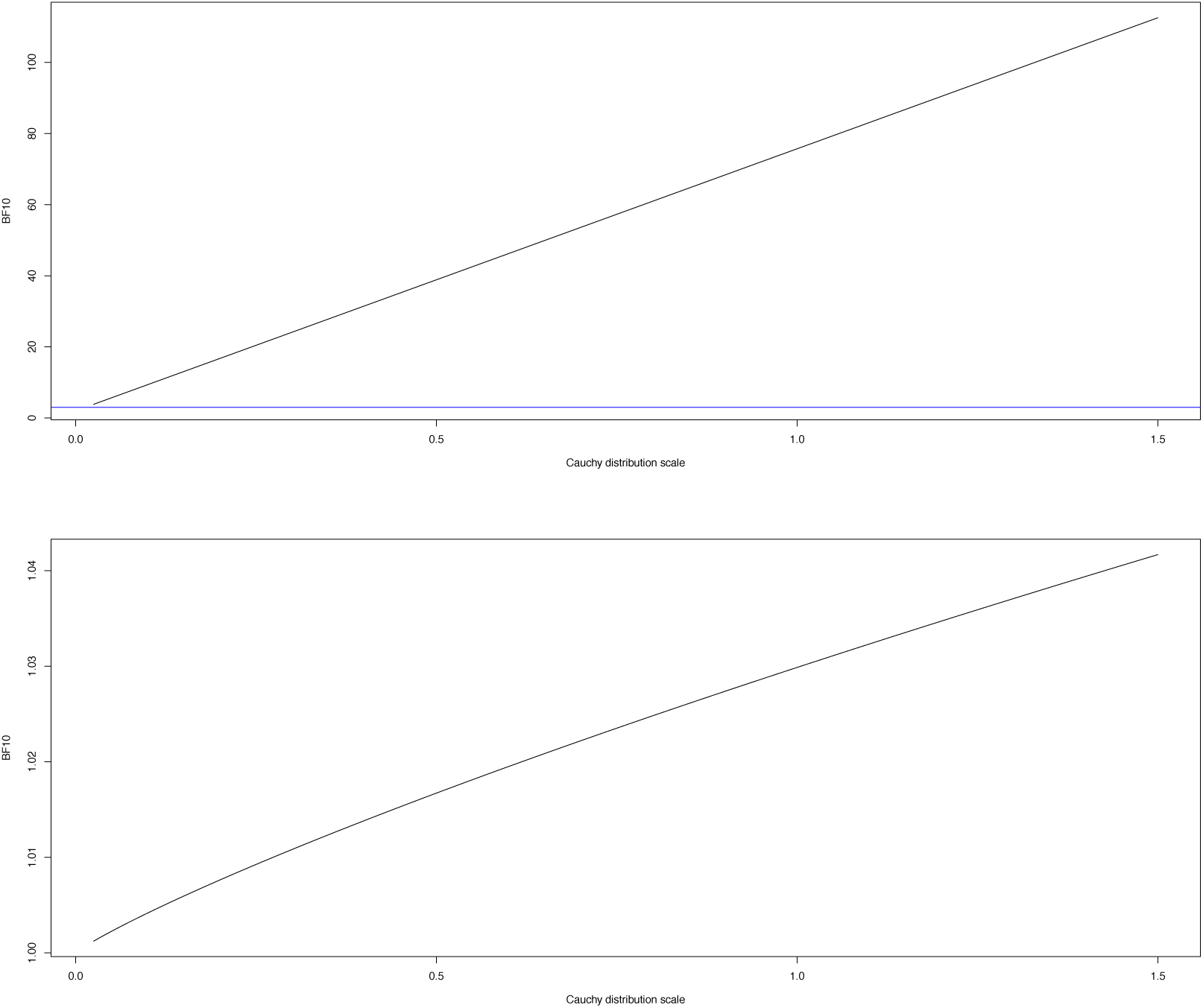
Changes in the Bayes Factor in two specified voxels according to changes in the Cauchy prior scale parameter (σ = .025 to 1.5). Top: change in the Bayes Factor in an active voxel, (45, 89, 36). Bottom: change in the Bayes Factor in an inactive voxel, (46, 64, 37). Black line: Bayes Factor. Blue line: Bayes Factor threshold, BF = 3.

**Figure S4.**
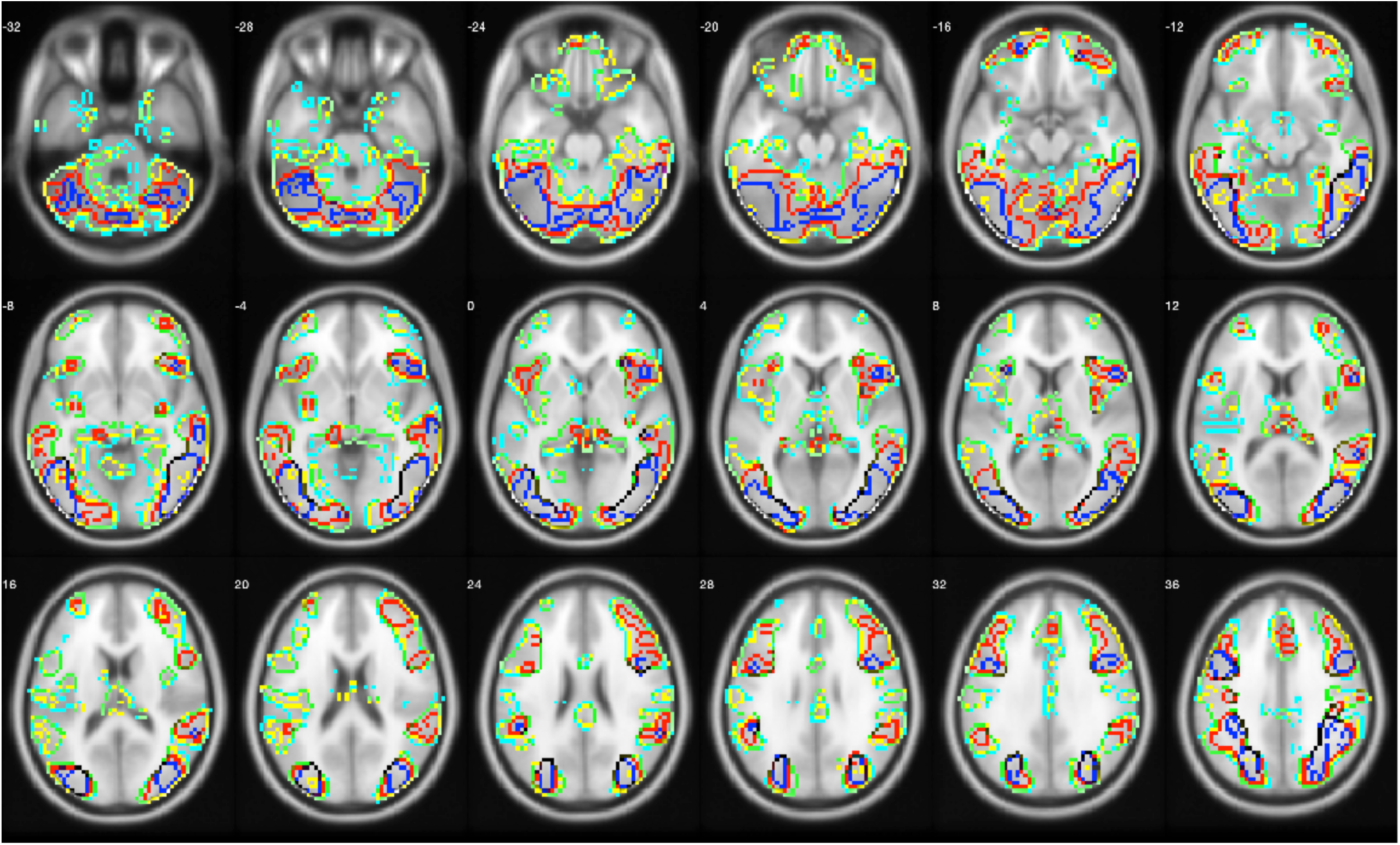
Results from the analysis of Amalric and Dehaene (2016). Red line: regions survived Bayesian thresholding with correction. Yellow line: regions survived Bayesian thresholding without correction. Blue line: regions survived classical thresholding with correction. Sky blue line: regions survived classical thresholding without correction.

**Figure S5.**
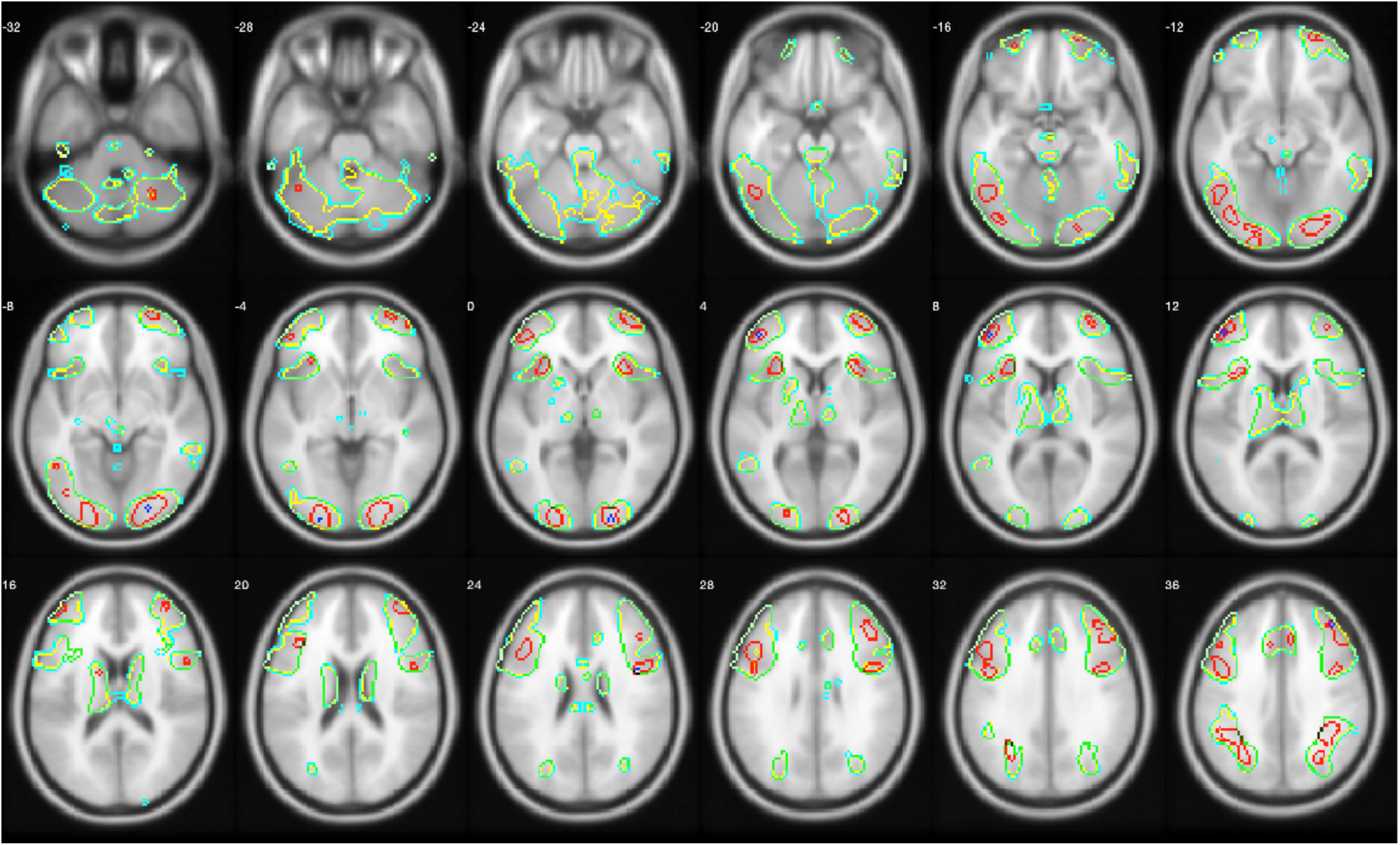
Results from the analysis of DeYoung et al. (2016). Red line: regions survived Bayesian thresholding with correction. Yellow line: regions survived Bayesian thresholding without correction. Blue line: regions survived classical thresholding with correction. Sky blue line: regions survived classical thresholding without correction.

**Figure S6.**
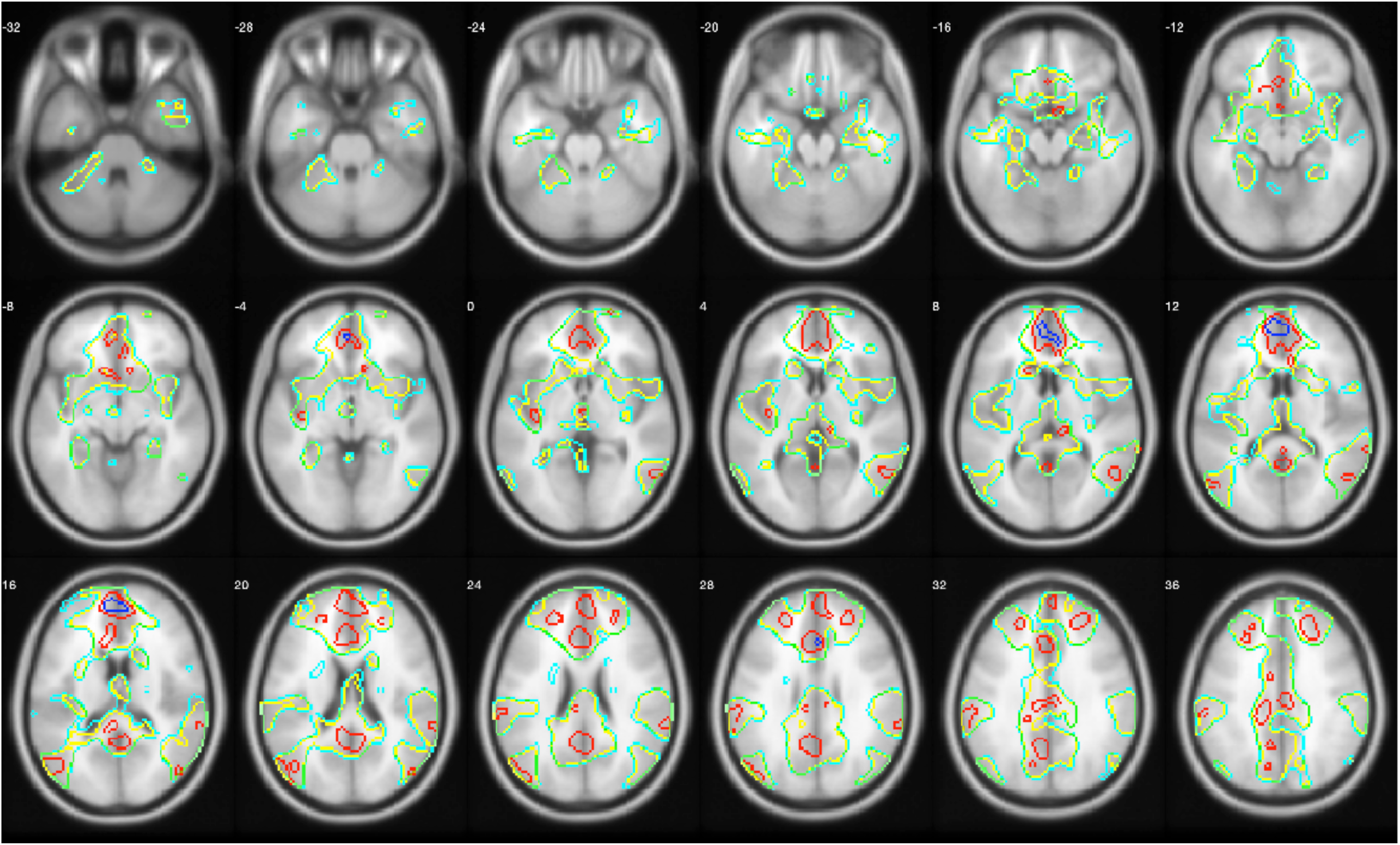
Results from the analysis of Han et al. (2016). Red line: regions survived Bayesian thresholding with correction. Yellow line: regions survived Bayesian thresholding without correction. Blue line: regions survived classical thresholding with correction. Sky blue line: regions survived classical thresholding without correction.

**Figure S7.**
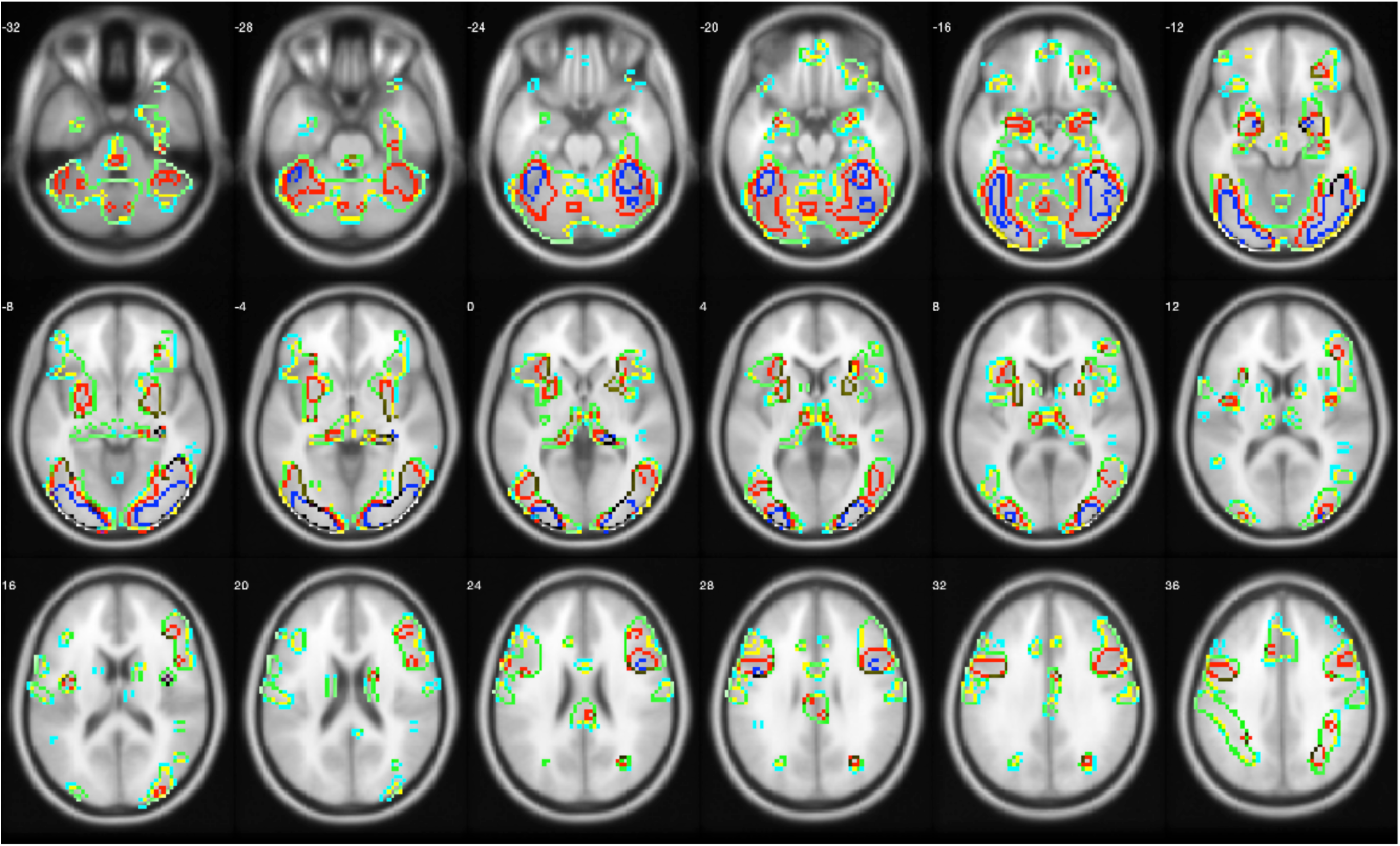
Results from the analysis of Henson et al. (2002). Red line: regions survived Bayesian thresholding with correction. Yellow line: regions survived Bayesian thresholding without correction. Blue line: regions survived classical thresholding with correction. Sky blue line: regions survived classical thresholding without correction.

**Figure S8.**
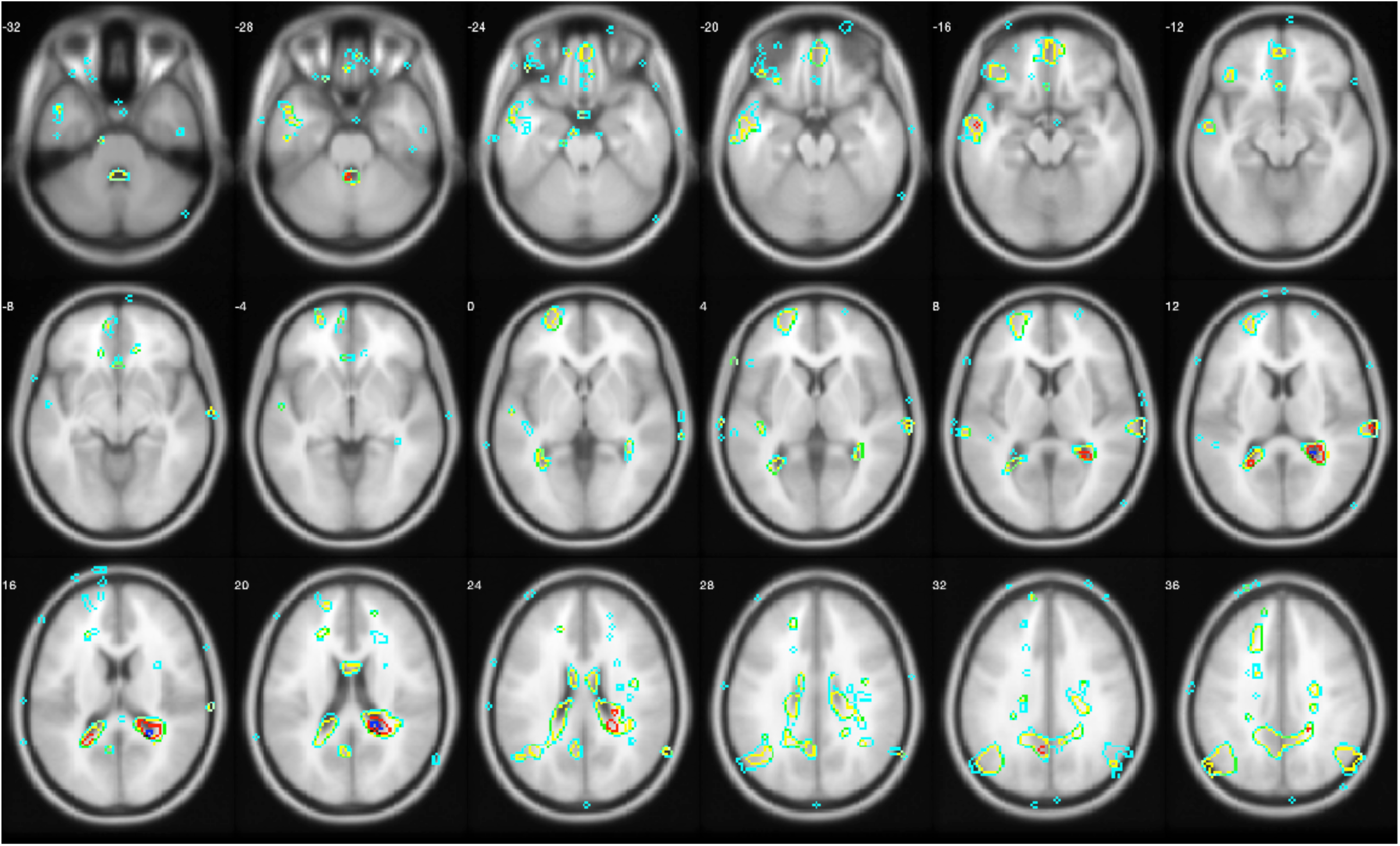
Results from the analysis of Kievit et al. (2016). Red line: regions survived Bayesian thresholding with correction. Yellow line: regions survived Bayesian thresholding without correction. Blue line: regions survived classical thresholding with correction. Sky blue line: regions survived classical thresholding without correction.

## Notes

https://osf.io/dbmu9/

